# Gene amplification during differentiation of mesenchymal stem cells towards chondrocytes

**DOI:** 10.64898/2026.06.16.732523

**Authors:** Paula E. Schwarz, Magali Cucchiarini, Shusruto Rishik, Andreas Keller, Eckart Meese, Ulrike Fischer

**Affiliations:** Department of Human Genetics, Saarland University, Building 60, 66421 Homburg/Saar, Germany; Center of Experimental Orthopedics, Saarland University Medical Center, 66421 Homburg/Saar, Germany; Chair of Clinical Bioinformatics, Saarland University, Saarbrücken, 66123, Germany

**Keywords:** gene amplification, CDK4, MDM2, chondrogenesis

## Abstract

For decades gene amplifications were described as an attribute of tumor cells and as a physiological mechanism to increase gene copy numbers for the higher protein demand during development of amphibians and flies. An increasing number of publications describe gene amplifications in normal mammalian cells during differentiation. Many amplified genes detected in tumor cells overlap with amplified genes detected during stem cell differentiation. Since stem cells have a valuable potency in regenerative therapies and since cartilage regeneration is a highly demanded therapeutic strategy, we investigated gene amplification dynamics during chondrogenic differentiation of human mesenchymal stem cells (hMSCs). Using quantitative PCR, we analyzed copy number changes for genes previously implicated in differentiation as well as genes amplified in chondrosarcoma including *CDK4, MDM2, AGAP2, CPT1B, SHANK3, TRIB1*, and *MYC*. Amplifications were transient and stage-specific: *CDK4, CPT1B*, and *SHANK3* exhibited the highest copy number increases at day 2, followed by a gradual decline by day 7, while *AGAP2* and *MDM2* increased later in differentiation. Laser microdissection of toluidine blue-stained areas revealed heterogeneity in amplification patterns: *CDK4* amplification was prominent in areas lacking or showing moderate proteoglycan deposition; *CPT1B* amplification occurred in regions with absent, moderate, or intense proteoglycan deposition; and *SHANK3* amplification was restricted to areas with intense proteoglycan deposition. Notably, regions with the strongest proteoglycan staining exhibited no gene amplification, suggesting that gene amplification is an early, transient event that diminishes as differentiation progresses. These findings highlight gene amplification as a mechanism during chondrogenesis, potentially critical for early differentiation stages and genome stability in mature cells.

## Introduction

Gene amplification is a genetic process where the copy number of certain genes increases, without a proportional increase of the genome. Gene amplification is not only a hallmark of genetic alterations in tumor cells but is also well described and studied since decades as physiological mechanism in amphibians and flies. For example, during the development of *Drosophila melanogaster* and *Xenopus laevis*, specific DNA regions are amplified within defined developmental time windows [1].

There is growing evidence that gene amplification plays a physiological role during differentiation of stem and progenitor cells. Previously we described gene amplification during differentiation of mesenchymal stem cells towards osteoblasts and adipocytes [2]. Interestingly, gene amplifications that arise during stem cell differentiation often overlap with those found in the corresponding tumor types. For example, the genes *CDK4* and *MDM2* were shown to undergo co-amplification during the differentiation of mesenchymal stem cells (MSCs) into adipocytes, mirroring the high-level amplifications characteristic of liposarcoma [3]. Similarly, several studies on chondrosarcoma have reported *MYC* gene amplification on chromosome 8, with array-CGH analyses demonstrating that *MYC* amplification occurs more frequently in higher-grade and dedifferentiated chondrosarcomas [4, 5]. In addition, genomic profiling of chondrosarcoma revealed several amplified chromosome regions including regions on chromosomes 5, 12, 19 and 22 [6]. Notably, amplification of *CDK4* without accompanying *MDM2* co-amplification has also been documented in at least one chondrosarcoma case, further underscoring the heterogeneity and potential clinical relevance of these genetic events [7].

Mesenchymal stem cells (MSCs) possess significant potential for therapeutic applications in tissue regeneration. One of the most ambitious investigations is probably regeneration of articular cartilage since many health restrictions are based on cartilage destruction [8-10]. Understanding both the regenerative capacity and the potential risks caused by genetic changes that are associated with directing hMSCs toward a chondrogenic lineage is therefore essential. Here we set out to investigate whether gene amplifications are detectable during chondrogenic differentiation of human mesenchymal stem cells.

## Materials and methods

### Cell culture and differentiation

Human mesenchymal stem cells (hMSC) from bone marrow were purchased form PromoCell (Heidelberg, Germany) were cultured until they form spheres in U-bottom 96-well plates or near confluent on Zeiss FrameSlides PET (polyethylene terephthalate) membrane for laser microdissection. Chondrocyte differentiation was induced in conditions using chondrogenic differentiation medium (PromoCell, Heidelberg, Germany) [11] for 5-7 days.

A detailed characterization of hMSCs is provided in the supplier’s certificates of analysis for Lot numbers 488Z010 (female donor, 79 years) and 479Z025 (male donor, 80 years), both of which were used in this study. As indicated in these certificates, surface marker analysis at passage 3 showed that 100% of cells were positive for CD73, CD90 and CD105, while 99-100% were negative for CD34 CD45. The certificates also document differentiation potential toward adipogenic, osteogenic, and chondrogenic lineages. All experiments were performed using cells at passage 3.

### RNA-Seq

RNA was isolated using miRNAeasy Kit (Qiagen, Hilden, Germany) and used for next generation sequencing. Sequencing was performed on an DNBSEQ-G400RS instrument by the NGS Sequencing Facility of Saarland University using the 100bp paired end sequencing strategy. Paired-end fastq files were processed using the mRNA module of SnakePipes 3.0.0 [12] The reads were aligned using STAR using default settings provided by snakepipes [13] against GRCh38 [14] in order to generate bam and bai files. Quality control checks were carried out using multiQC [15].

### Toluidine Blue staining of proteoglycan during differentiation

HMSCs were cultivated and differentiated as described above. For toluidine blue staining cells were washed once with PBSx1 for 10 minutes and either fixed with 4% PFA (paraformaldehyde) for 5 minutes or with ice cold methanol for 10 minutes. Toluidine staining solution (10% Toluidine Blue, 1% Borax) was applied for 15 minutes, and excess staining was removed by three PBSx1 washing steps [16]. Differentiation was checked with a bright-field microscope at 20x magnification.

### Laser microdissection of differentially toluidine-blue stained areas

For laser microdissection hMSCs were differentiated for 5 days towards chondrocytes and stained with toluidine blue for proteoglycan detection of differentiating cells. In total 20 areas (for each staining subpopulation: grey, light blue, blue and dark blue) of 0.76 mm^2^ were laser microdissected, using the PALM^®^ MicroBeam Microdissection System (Carl Zeiss AG). These experiments were done in three biological replicates. DNA extraction was performed using the QIAamp DNA Micro Kit (Qiagen) according to the manufacturer’s instructions. The final eluate was concentrated to a total volume of approximately 12 µl and DNA concentration was determined using Qubit™ dsDNA HS Assay Kit (Thermo Fisher Scientific, Waltham, USA).

### qPCR analysis

TaqMan Copy Number Assays for genes *CDK4* (Hs00957586_cn), *MDM2* (Hs00181272_cn), *AGAP2* (Hs01425078_cn), *CPT1B* (Hs01741692cn), *MYC* (Hs02602824_cn), *SHANK3* (Hs04081743_cn), and *TRIB1* (Hs00836770_cn) were performed following manufacturer’s instructions (Applied Biosystems®, Pleasanton, USA). *RNaseP* TaqMan Copy number reference assay was used for relative quantitation of copy number of target genes. DNA from not differentiated controls of hMSCs cultivated in DMEM was used as control standard for normal diploid copy number. TaqMan assays were run in four technical replicates and results were analyzed using StepOne™ Software v2.0 and CopyCaller™ software (Applied Biosystems® by Life Technologies Corporation) for differentiation in U-bottom 96-well plates. TaqMan Copy Number Assays for genes *CDK4* (Hs00957586_cn), *AGAP2* (Hs01425078_cn), *CPT1B* (Hs01741692cn), *SHANK3* (Hs04081743_cn), and *TRIB1* (Hs00836770_cn) were performed with 250 pg DNA per reaction. TaqMan assays were run in two technical replicates and results were analyzed using StepOne™ Software v2.0 and CopyCaller™ software for laser microdissected areas. For analysis of this low amount of DNA input the Ct-threshold was set to exclude values above 35.

Analyses were conducted using three biological replicates, each measured in two technical replicates. Statistical significance was assessed using an unpaired *t*-test (Microsoft Excel).

## Results

HMSCs were differentiated in U-bottom 96-wells and differentiation was confirmed by toluidine blue staining and RNA-Seq analysis of chondrogenic differentiation specific genes *SOX9, COL2A1* and *ACAN [17, 18]*. Already at 5 d post differentiation induction cells revealed three-dimensional cell growth as spheres and high intensity of proteoglycan specific toluidine blue staining (Figure 1). Expression analysis of two independent biological replicates revealed an increase of *SOX9* expression after 2 days and a further increase after 7 days of differentiation and an increase of *COL2A1* and *ACAN* expression after 5days and 7 days of differentiation (Figure 2). To detect gene amplifications, we analyzed bulk DNA from hMSCs differentiated toward chondrocytes for 1, 2, 5, and 7 days, as well as from undifferentiated hMSCs serving as controls. Amplification was assessed by quantitative PCR (TaqMan) in four replicates, and data were evaluated using the CopyCaller software (Applied Biosystems).

**Figure 1.**
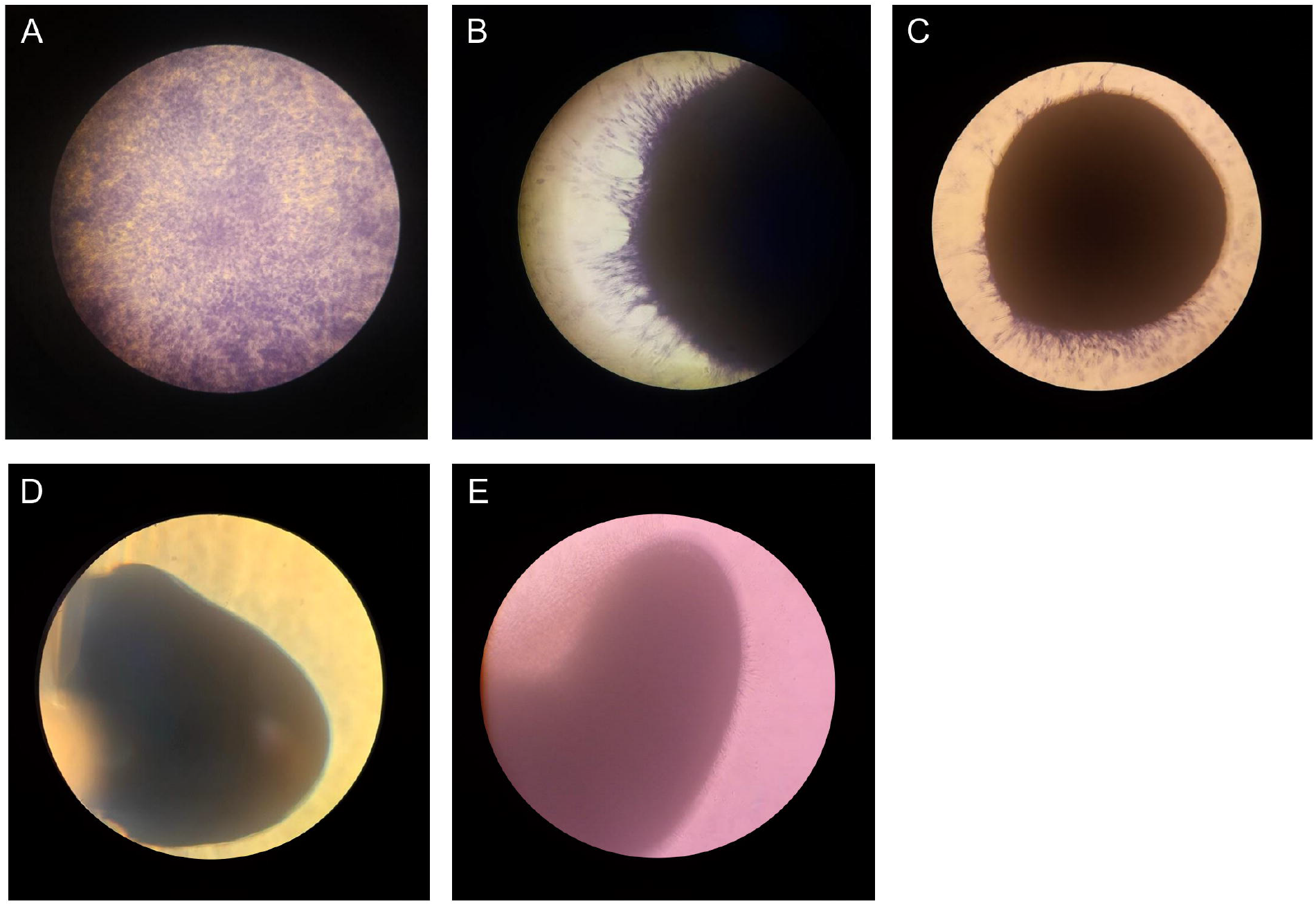
Proteoglycan and spheroid formation during chondrocyte differentiation. HMSCs were differentiated in U-bottom wells, and spheroid formation together with proteoglycan deposition using toluidine blue staining was monitored over 14 days. Proteoglycan staining by toluidine blue was detectable as early as day 2 in the absence of spheroid formation (A). Spheroid formation became evident by day 5 (B). Strong toluidine blue staining was observed in all spheroids at days 7 (C) and 14 (D). An unstained spheroid at day 14 is shown as a reference (E). Images were taken at 20x magnification.

**Figure 2.**
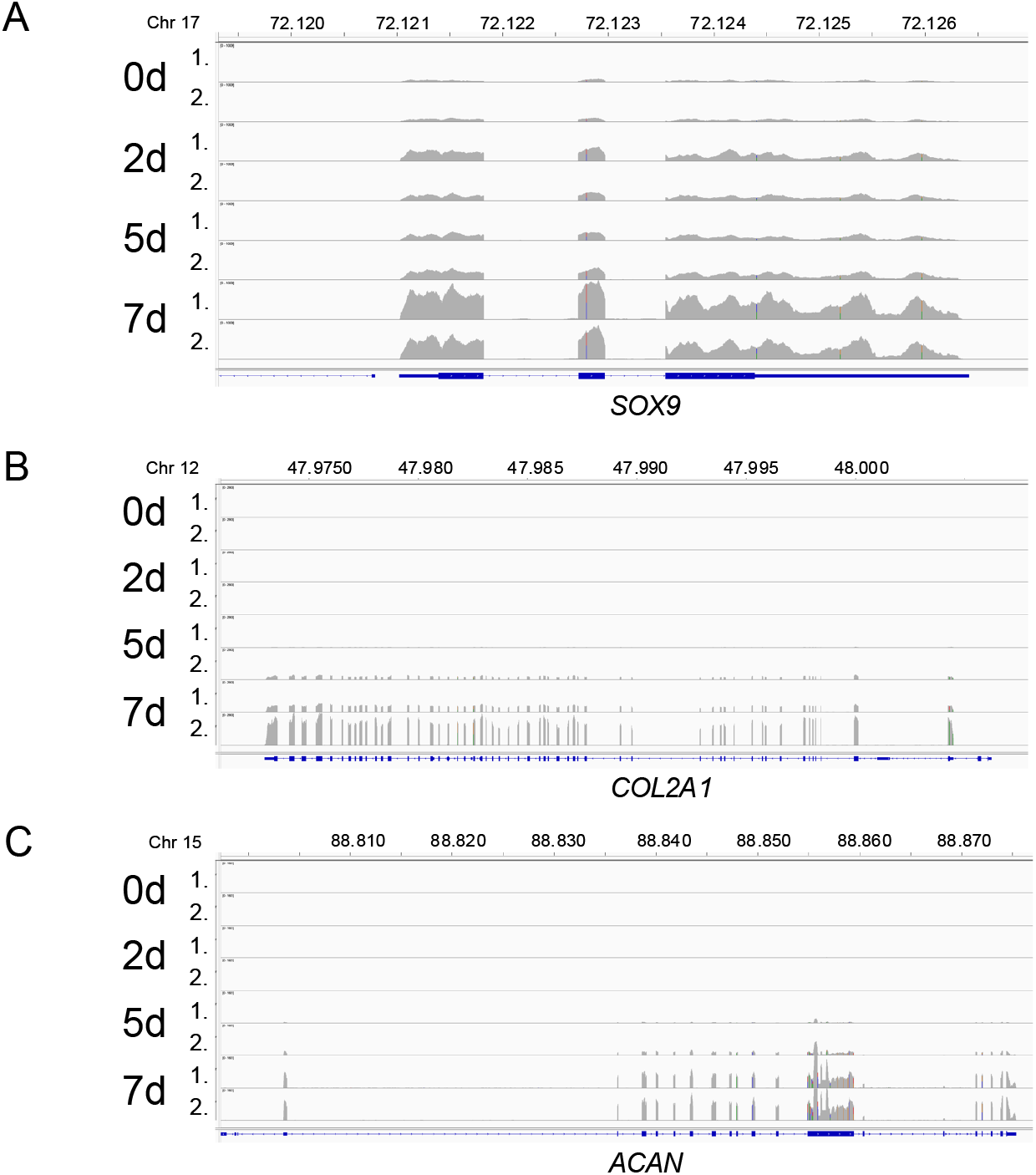
Expression of *SOX9, COL2A*, and *ACAN* during chondrocyte differentiation. RNA-seq data from two biological replicates (shown as 1. and 2. on the left) are visualized using IGV (Integrative Genomics Viewer). Undifferentiated hMSCs exhibited minimal *SOX9* expression; levels increased at days 2 and 5 and reached a maximum at day 7, consistent with chondrocyte differentiation (A). *COL2A1* (B) and *ACAN* (C) expression were undetectable at day 2 but became apparent at day 5 and peaked at day 7, further confirming successful chondrogenic differentiation. X-axis indicates chromosome localization in Mb and y-axis RNA-seq results represented as read-height.

For our investigation we selected genes previously found to be amplified during osteoblast and adipocyte differentiation and genes that were known to be amplified in tumors derived from chondrocytes including genes on chromosome 12, chromosome 22 and chromosome 8 [2, 6]. Copy number was determined for genes *AGAP2, CDK4* and *MDM2* localized on chromosome 12, *CPT1B* and *SHANK3* localized on chromosome 22 and *TRIB1* and *MYC* localized on chromosome 8. As shown in Figure 3 after 1d of differentiation none of the investigated genes revealed a copy number increase. After 2d of differentiation *CDK4, CPT1B* and *SHANK 3* revealed a 60% copy number increase, *AGAP2* and *TRIB1* exhibited a 20% copy number increase; *MDM2* and *MYC* remained unchanged (Figure 3). After 5d of differentiation *AGAP2* and *MDM2* displayed increases, whereas *CDK4, CPT1B* and *SHANK3*, which were highest at 2 days, decreased and *MYC* still remained unchanged. After 7d of differentiation *CDK4, CPT1B, SHANK3*, and *TRIB1* decreased further, *AGAP2* increased, and no further copy number change was detected for *MDM2. MYC* showed no change throughout the 7-day differentiation period.

**Figure 3.**
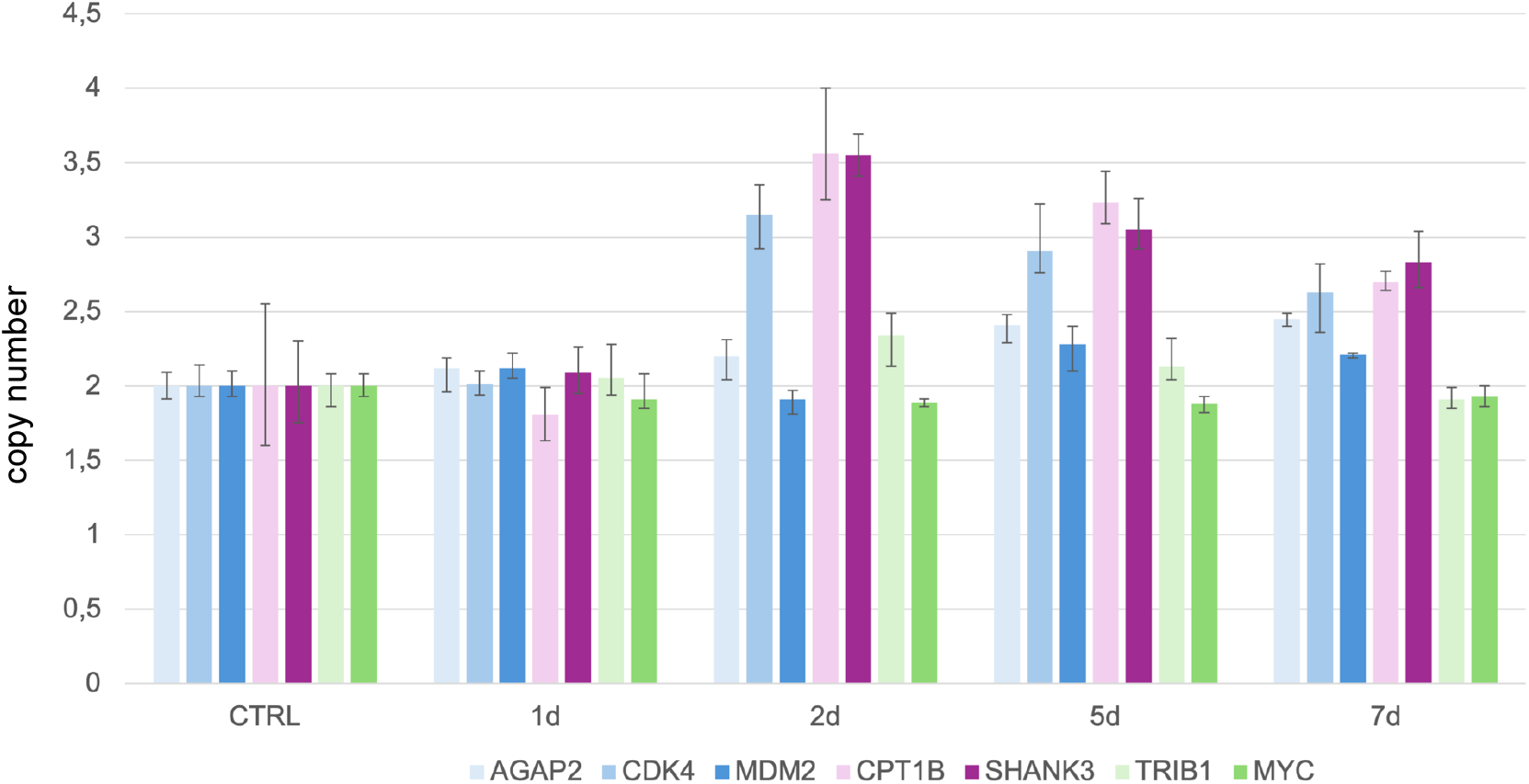
Amplification analysis of *AGAP2, CDK4, MDM2, CPT1B, SHANK3, TRIB1* and *MYC* using qPCR during chondrogenic differentiation. Amplification of *AGAP2, CDK4, MDM2, CPT1B, SHANK3, TRIB1* and *MYC* was analyzed by qPCR using TaqMan copy number assays and *RNAseP* was used as reference TaqMan assay. DNA from undifferentiated hMSCs (CTRL) served as standard for normal diploid copy number with copy number values indicated on the y-axis and normal diploid copy number is referred as 2. Copy numbers are shown as mean from four technical replicates with vertical lines indicating the range. Chondrogenic-induced hMSC cells were analyzed at four time points after differentiation induction (1d, 2d, 5d and 7d). *CPT1B* and *SHANK3* revealed the highest copy numbers of all investigations after 2d of differentiation and a decrease until 7d. *CDK4* revealed also a high copy number after 2d of differentiation induction (green) and a decrease until 7d. *TRIB1* revealed a copy number increase only 2d after differentiation induction. *AGAP2* and *MDM2* revealed a copy number increase only after 5 days and 7 days. *MYC* revealed no copy number increase at any time point after differentiation induction.

Previous published investigations on gene amplification during differentiation revealed a high heterogeneity of gene amplification between single cells. During osteoblast differentiation for example, cells with high level amplification were detected next to cells with normal copy number and during neural differentiation composition and copy number of co-amplified genes varied between neighboring cells [2, 19]. One possible reason for the differences in gene copy numbers is differences in the degree of differentiation upon differentiation induction. After five days upon differentiation induction, we performed laser microdissection on hMSCs after toluidine blue staining for detection of proteoglycan production that is indicative for chondrocyte differentiation (Figure 4 A-D). Toluidine blue stain revealed heterogenous blue intensities with areas of grey (A), light blue (B), blue (C) and dark blue (D) staining as result of different amounts of produced proteoglycans indicating different stages of differentiation.

**Figure 4.**
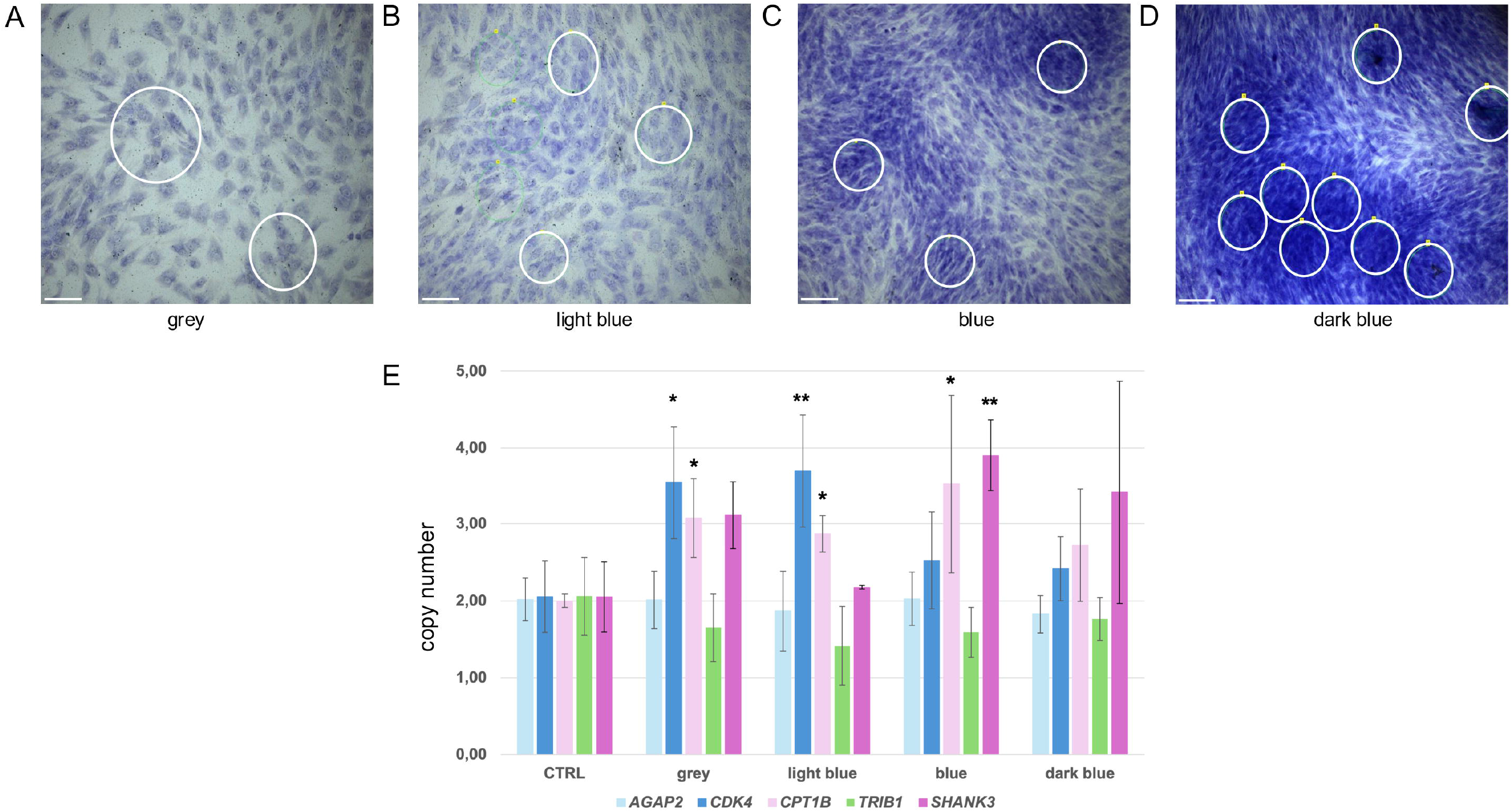
Amplification analysis of *AGAP2, CDK4, CPT1B, SHANK3* and *TRIB1* after laser microdissection. HMSCs were differentiated towards chondrocytes for 5d on PET frame slides for laser microdissection. Cells were seeded at high density and slide was stained with toluidine blue for proteoglycan detection during differentiation. Different areas of staining intensity could be detected and were defined as areas with grey cells (A), areas with light blue cells (B), areas with blue cells (C) and areas with dark blues cells (D), all indicating stages during the differentiation towards chondrocytes. In total for each area type twice 20 areas were laser microdissected and used for DNA isolation. Experiments were done in three biological replicates. Examples of laser microdissected areas are circled in white and scale bars represent 150µm. Results of amplification analysis of our qPCR using TaqMan copy number assays and *RNAseP* as reference are shown as diagram with *AGAP2, CDK4* and *CPT1B SHANK3* and *TRIB1* (E). DNA from undifferentiated hMSCs (CTRL) served as standard for normal diploid copy number with copy number values indicated on the y-axis and normal diploid copy number is referred as 2. Copy numbers are presented as means of two technical replicates across three biological replicates, with error bars representing standard deviation. Statistical significance is indicated as *p* < 0.05 (*) and *p* < 0.01 (**). *CDK4* revealed the highest copy number in areas with grey and light blue cells indicating gene amplification at the beginning of differentiation. *CPT1B* revealed high copy numbers in grey, light blue and blue cells indicating gene amplification during differentiation and *SHANK3* revealed the highest copy numbers in areas with blues cells indicating gene amplifications during later differentiation progression. *AGAP2* and *TRIB1* revealed no copy number increase. In areas with dark blues cells no significant amplification was detectable indicating a reduction of gene amplification towards final differentiation.

In total we laser microdissected 20 areas for each blue intensity as shown in Figure 4 A-D. Although the laser microdissected regions were comparable in size the amount of isolated DNA varied between 1.8 ng for grey and light blue areas and 6 ng for blue and dark blue areas. Copy analysis was done by TaqMan qPCR with 250 pg laser microdissected DNA for each experiment. Analyses were performed using three biological replicates, each with two technical replicates, and undifferentiated hMSC DNA served as the control. Copy number variations for *AGAP2, CDK4, CPT1B, TRIB1*, and *SHANK3* are shown in Figure 4E. A significantly increased *CDK4* copy number was observed in cells from grey and light blue regions, corresponding to absent or low proteoglycan content. *CPT1B* copy number was significantly elevated in grey, light blue, and blue regions, indicative of absent to early proteoglycan deposition. In contrast, a significant increase in SHANK3 copy number was detected in blue regions, reflecting higher proteoglycan levels. Interestingly, dark blue areas with highest proteoglycan levels revealed no significant copy number increase for the investigated genes. *AGAP2* showed no copy number increase in the laser microdissected probes for all staining intensities. TaqMan analysis of *MDM2* revealed no results for technical reasons.

## Discussion

Gene amplification has long been associated with tumorigenesis and multidrug resistance, although its physiological role was first recognized in *Drosophila melanogaster* during eggshell development [20, 21]. Here, we identified gene amplifications in hMSCs during chondrogenic differentiation. Previous fluorescence-*in-situ*-hybridization (FISH) analyses of hMSCs differentiating into adipocytes and osteoblasts revealed heterogenous populations, with amplified and non-amplified cells coexisting [2]. However, FISH analysis proved challenging for chondrocyte differentiation due to the three-dimensional growth pattern and proteoglycan production, which interfere with FISH. Here, we investigated gene amplification during the first seven days of chondrocyte differentiation using qPCR and Taq Man assays targeting genes previously shown to be amplified during osteoblast and adipocyte differentiation, as well as genes that were known to be amplified in chondrosarcoma [2, 6].

DNA sequence amplification, as defined by Bostock CJ in 1986, refers to any event that increases a gene’s copy number per haploid genome above the level characteristic for that organism [1]. In accordance with a criterion defined in our recent study [22], qPCR-derived copy number values exceeding 2.3 were considered indicative of gene amplification. To distinguish gene amplification from whole-chromosome duplication, we included genes located on the same chromosomes (*AGAP2, CDK4*, and *MDM2* on chromosome 12; *CPT1B* and *SHANK3* on chromosome 22; and *TRIB1* and *MYC* on chromosome 8).

During chondrocyte differentiation, qPCR analysis revealed amplification of several investigated genes at days 2 and 5, followed by a decline by day 7. To assess stage-specific amplification, we performed laser microdissection of toluidine blue-stained areas after five days of differentiation, when proteoglycan deposition was evident. Cells were grown on PET membrane slides, enabling precise isolation of regions with varying staining intensities.

Amplification patterns differed across differentiation stages: *CDK4* showed the highest copy number in regions lacking and low proteoglycans, indicating early activation, whereas *SHANK3* peaked in areas with strong proteoglycan staining, suggesting stage-specific amplification. Regions with the most intense staining displayed no significant amplifications for all tested genes, consistent with cessation of amplification during terminal differentiation. Although *AGAP2* increased in bulk DNA at days 5 and 7, it was undetectable in microdissected samples, and *MDM2* yielded no results, likely due to technical limitations of microdissection and DNA isolation. A possible reason for failure of *MDM2* qPCR amplification could be the very low amount of DNA per reaction (250 pg) and degradation effects during laser microdissection and DNA isolation. In this respect we would like to mention that gene amplification analysis using microdissected samples is a starting point to address gene amplifications in a heterogenous cell population but has limitations because of low DNA yield and degradation during laser exposure. Furthermore, although *CPT1B* and *SHANK3* are both located on chromosome 22, their distinct amplification patterns across microdissected regions argues against whole-chromosome duplication and instead support gene-specific amplification events.

## Conclusion

In summary, we observed in both bulk and microdissected analyses that gene amplification is an early, transient process during chondrogenic differentiation, diminishing as cells mature— likely to preserve genomic stability. Since this study has limitations, further orthogonal validation methods including single cell analysis were needed. Future studies should explore mechanisms that regulate amplification cessation and their implications for regenerative therapies. Nevertheless, the current work provides valuable insights into advantages and capabilities including risks of these cells.

## Author contribution

P.E.S.: investigation, writing – review and editing; M.C.: investigation, writing – review and editing S.R.: formal analysis, data curation, writing – review and editing; A.K.: funding acquisition, writing – review and editing, E.M.: conceptualization, funding acquisition, writing – original draft, review and editing, U.F.: conceptualization, investigation, visualization, writing – original draft, review and editing.

## Acknowledgement

We appreciate the help of the *NGS Sequencing Facility* of Saarland University for providing support.

